# A Symmetric Systemic Challenge Elicits a Right-Biased Response Mediated by Vasopressin Signaling

**DOI:** 10.64898/2026.03.02.708998

**Authors:** Hiroyuki Watanabe, Yaromir Kobikov, Sara Yusuf Mohamed, Karen Rich, Daniil Sarkisyan, Olga Nosova, Alfhild Grönbladh, Mathias Hallberg, Jens Schouenborg, Georgy Bakalkin, Mengliang Zhang

**Affiliations:** Department of Pharmaceutical Biosciences, Uppsala University, SE-751 24 Uppsala Sweden; Department of Molecular Medicine, University of Southern Denmark, DK-5230, Odense M, Denmark; Department of Immunology, Genetics and Pathology and Science for Life Laboratory, Uppsala University, SE-751 05 Uppsala, Sweden; Neuronano Research Center, Department of Experimental Medical Science, Lund University, 223 63 Lund, Sweden

**Keywords:** water deprivation, operational asymmetry, vasopressin, hindlimb postural asymmetry, symmetric systemic challenge, left-right balance

## Abstract

Bilaterian animals exhibit *operational* (functional) asymmetry—population-level, directional left–right differences in physiology and behavior, including responses to spatially symmetric environmental challenges. Whether such symmetry-to-asymmetry conversion can be driven at the systems level by neurohormonal regulators remains unclear. Here we tested whether a spatially symmetric neuroendocrine challenge—water deprivation (WD)—can elicit a directional left–right physiological response in rats using hindlimb postural asymmetry (HL-PA), a binary readout that quantifies left- versus right-sided hindlimb flexion.

Twenty-four hours of WD induced robust HL-PA with right hindlimb flexion, revealed under anesthesia. The asymmetry persisted after complete thoracic spinal cord transection, suggesting that humoral signaling, rather than descending neural commands, may maintain the postural bias. Because dehydration recruits the hypothalamic–neurohypophysial arginine vasopressin (AVP) system, we next tested AVP receptor involvement. Both a V1B antagonist (SSR-149415) and a V1A/V2 antagonist (conivaptan) abolished WD-induced HL-PA, supporting an AVP-dependent mechanism that likely operates at least two anatomical sites. AVP signaling may involve pituitary V1B–dependent endocrine output and spinal V1A actions; consistent with the latter, expression of AVP V1A receptors is right-biased in lumbar spinal cord.

Together, these findings identify WD as a symmetric systemic challenge capable of imposing a directional peripheral set-point, and implicate vasopressin signaling in symmetry breaking and left–right physiological regulation.

**Visual summary:** 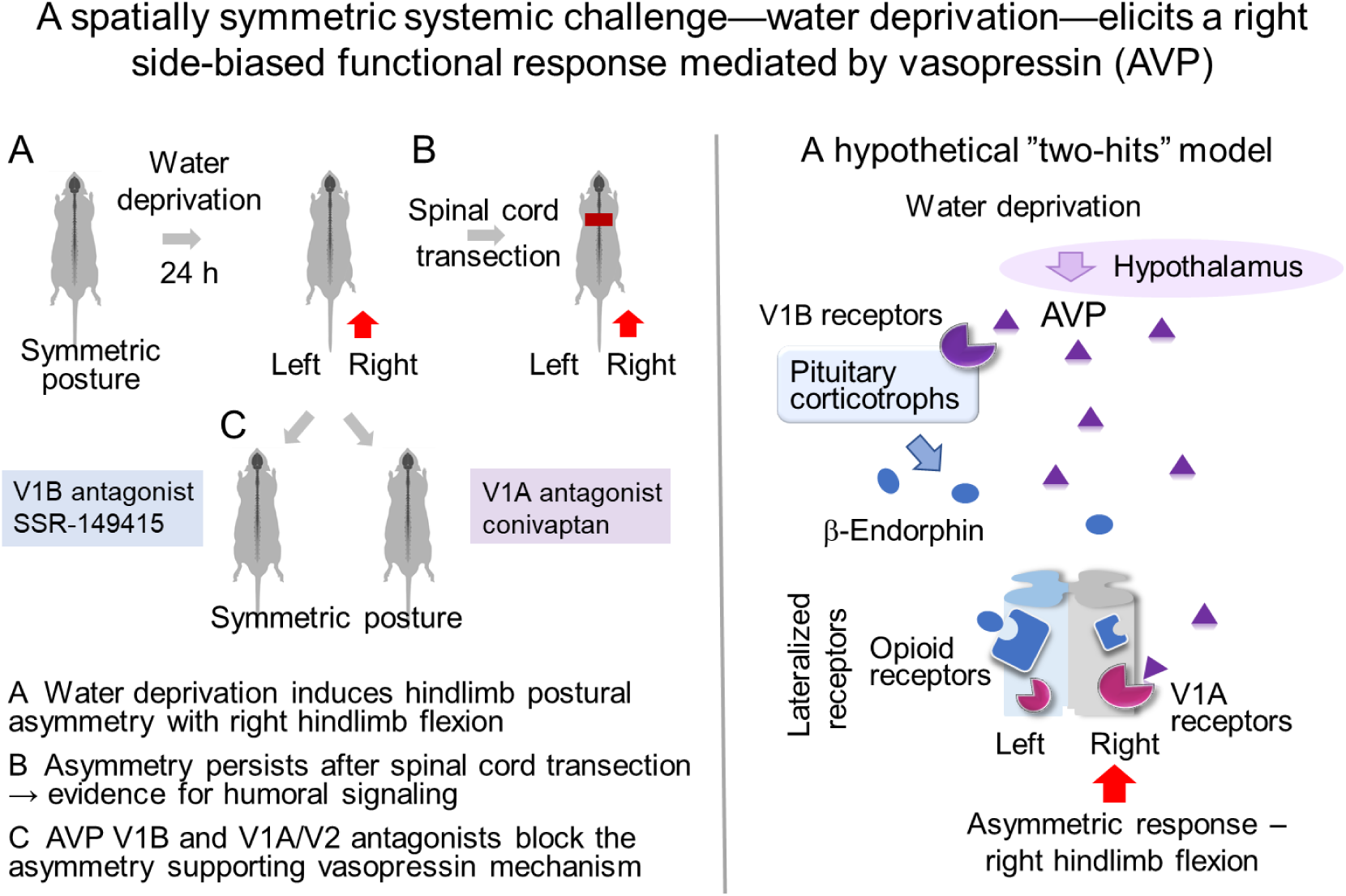

## Introduction

Alongside developmental left–right asymmetry (e.g., visceral laterality established during embryogenesis) [1,2], bilaterian animals also exhibit operational (functional) asymmetry—population-level, directional left–right differences in physiological or behavioral processes that emerge during activation despite broadly symmetric anatomy and symmetric stimulation [3,4]. Here, “symmetric” denotes inputs that carry no inherent left–right information; operational asymmetry is used as an umbrella term that includes, but is not restricted to, hemispheric specialization.

Operational asymmetry is conserved across taxa. In adult *C. elegans*, whole-animal salt changes are encoded asymmetrically by paired gustatory neurons (ASEL preferentially signals increases; ASER decreases), enabling directional chemotaxis in a symmetric stimulus field [5]. Similar population-level lateralization is observed in other bilaterians, including insects and vertebrates[3,4]. These observations support the general principle that bilaterian systems can transform symmetric inputs into asymmetric outputs via lateralized internal processing (“symmetric stimulus → symmetric gross anatomy → asymmetric response”).

A key mechanistic question is whether this directional conversion can be implemented or gated by systemic regulators, especially neuropeptides and protein hormones, which are particularly recruited by symmetric homeostatic and stress challenges [6]. In *C. elegans*, peptide hormone signaling reshapes laterality under symmetric conditions: ASE-derived INS-6 insulin signaling remodels salt-circuit computations [7], associative learning restructures circuit laterality through asymmetric insulin signaling and new synapse formation [8], and CAPA-1/NMUR-1 (NMU-like) neuropeptide signaling regulates expression of the learned lateralized state [9]. In turtles, opioid peptides acting via KOR and DOR which are enriched in the left and right visual cortices, respectively, modulate contralateral visual input [10,11]. In rodents, neuropeptides and neurohormones can modulate activity of lateralized circuits, and induce side-specific responses upon their symmetric administration [6]. Signaling by neuropeptides dynorphin, calcitonin gene-related peptide and substance P is lateralized in the parabrachial-to-amygdala circuit, which underlies the asymmetric processing of pain in mice [12–16]. The oxytocin control of auditory processing during maternal behavior is lateralized to the left versus right auditory cortex [17]. In rodents, ghrelin asymmetrically modulates brain metabolism, and CCK-8 produces opposite behavioral responses when microinjected into the left or right hemisphere [18,19].

Brain injury studies reveal the side-specific functions of arginine vasopressin (AVP) and opioid peptides as systemic mediators of contralateral injury effects on posture and the motor system [20–25]. Lesions in the left sensorimotor cortex produce hindlimb postural asymmetry with right-side limb flexion that is blocked by the AVP V1B receptor antagonist SSR-149415. Intravenous AVP mimics this contralateral effect, inducing HL-PA in animals with intact brains.

It is essential to clarify whether symmetric systemic challenges to the neuroendocrine system can produce population-level, directional left-right physiological and behavioral outputs and whether neurohormones with side-specific actions, such as AVP, can contribute to or regulate such transformations from symmetry to asymmetry. We therefore tested whether a systemic neuroendocrine challenge—water deprivation—can elicit HL-PA as a directional left–right response, and whether this response is mediated by the AVP mechanism. Dehydration lacks directional cues—left–right bias would argue for internally generated, neuroendocrine symmetry breaking—and recruits the hypothalamic–neurohypophysial AVP system [26–28]. As a readout for lateralized endocrine responses, we employed the HL-PA model—a binary outcome system that quantifies left- versus right-sided postural changes [21,22]. To isolate neuroendocrine signaling from descending neural input, the spinal cords were completely transected at the thoracic level in subset of the experiments.

## Results

### Water deprivation effects on hindlimb posture

Figures 1–2 present posterior summaries from Bayesian regression across experimental groups: posterior medians (black circles), 95% highest posterior density credible intervals (95% HPD; black horizontal lines), and posterior distributions (colored density plots). The 95% HPD interval denotes the range that contains 95% of the posterior probability mass for the parameter, and can be viewed as a Bayesian analogue of a confidence interval. Asymmetry was considered supported when the 95% HPD interval excluded zero and the corresponding multiplicity-adjusted P value was ≤ 0.05. These adjusted P values were produced in emmeans by applying its frequentist summary to the posterior mean and covariance implied by the Bayesian model estimates, with multiple-comparisons adjustment. We do not combine Bayesian and frequentist decision rules: credible intervals are treated as the primary inferential output, whereas adjusted P values are reported as supplementary, model-based summaries.

**Figure 1.**
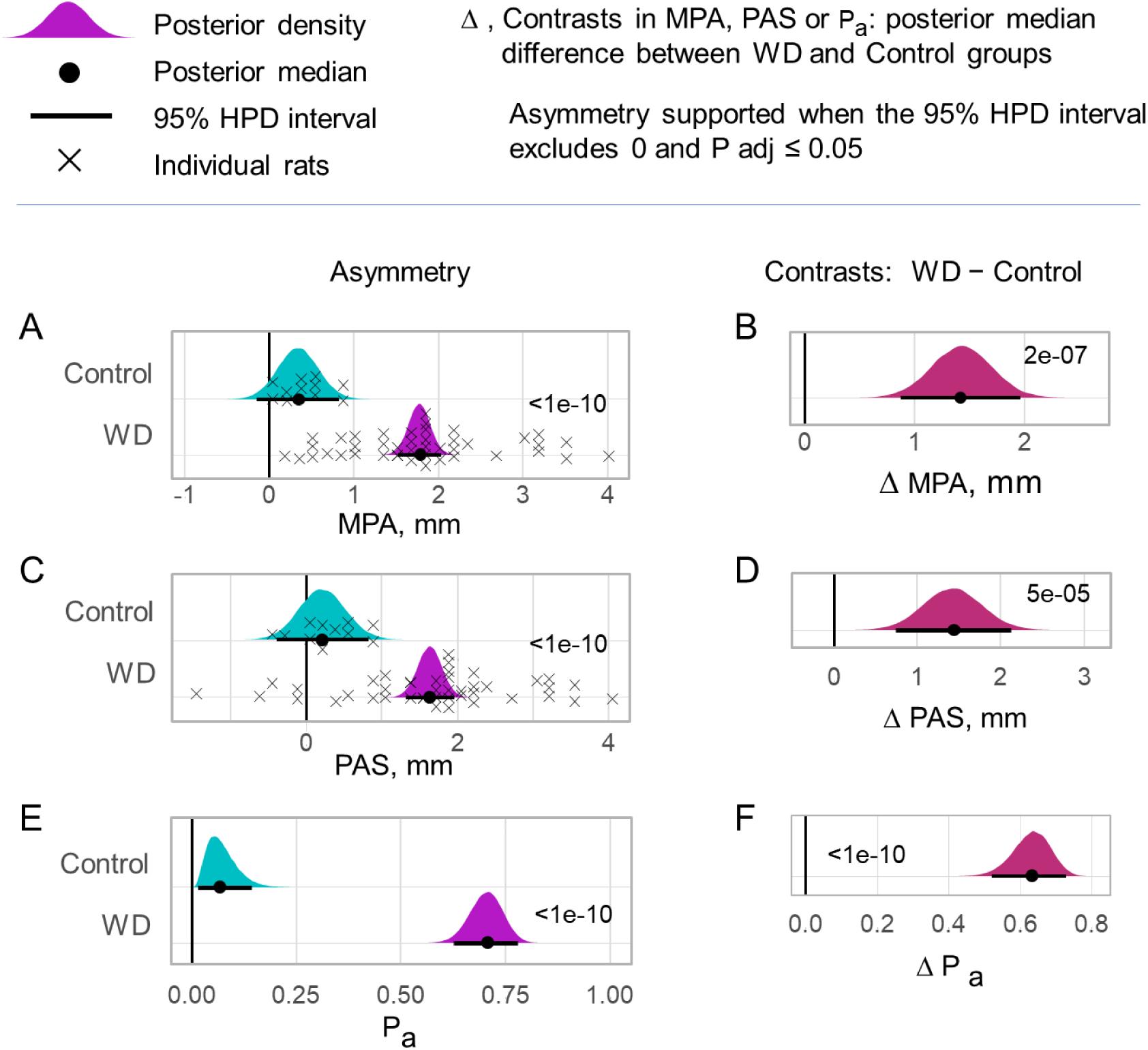
Water deprivation–induced hindlimb postural asymmetry (HL-PA). Magnitude of postural asymmetry (MPA), postural asymmetry score (PAS), and the proportion of asymmetric animals (P_a_) were quantified. PAS encodes directionality: negative values indicate left hindlimb flexion and positive values indicate right hindlimb flexion. **(A,C,E)** HL-PA in control (n = 12) and water-deprived (n = 43) rats; crosses denote individual animals. **(B,D,F)** Between-group contrasts (water-deprived minus control) for MPA (ΔMPA), PAS (ΔPAS), and Pa (ΔP_a_). Posterior summaries from Bayesian regression are shown as posterior medians (black circles), 95% highest posterior density (HPD) intervals (black horizontal lines), and posterior distributions (colored density plots). Effects were considered supported when the 95% HPD interval excluded zero (MPA ≠ 0, PAS ≠ 0 and P_a_ ≠ 0 in **A,C,E;** ΔMPA ≠ 0, ΔPAS ≠ 0 and ΔP_a_ ≠ 0 in **B,D,F**) and the corresponding multiplicity-adjusted P value was ≤ 0.05 (adjusted P values are shown on the plots). Credible intervals are the primary inferential summaries; adjusted P values are reported as supplementary model-based estimates.

**Figure 2.**
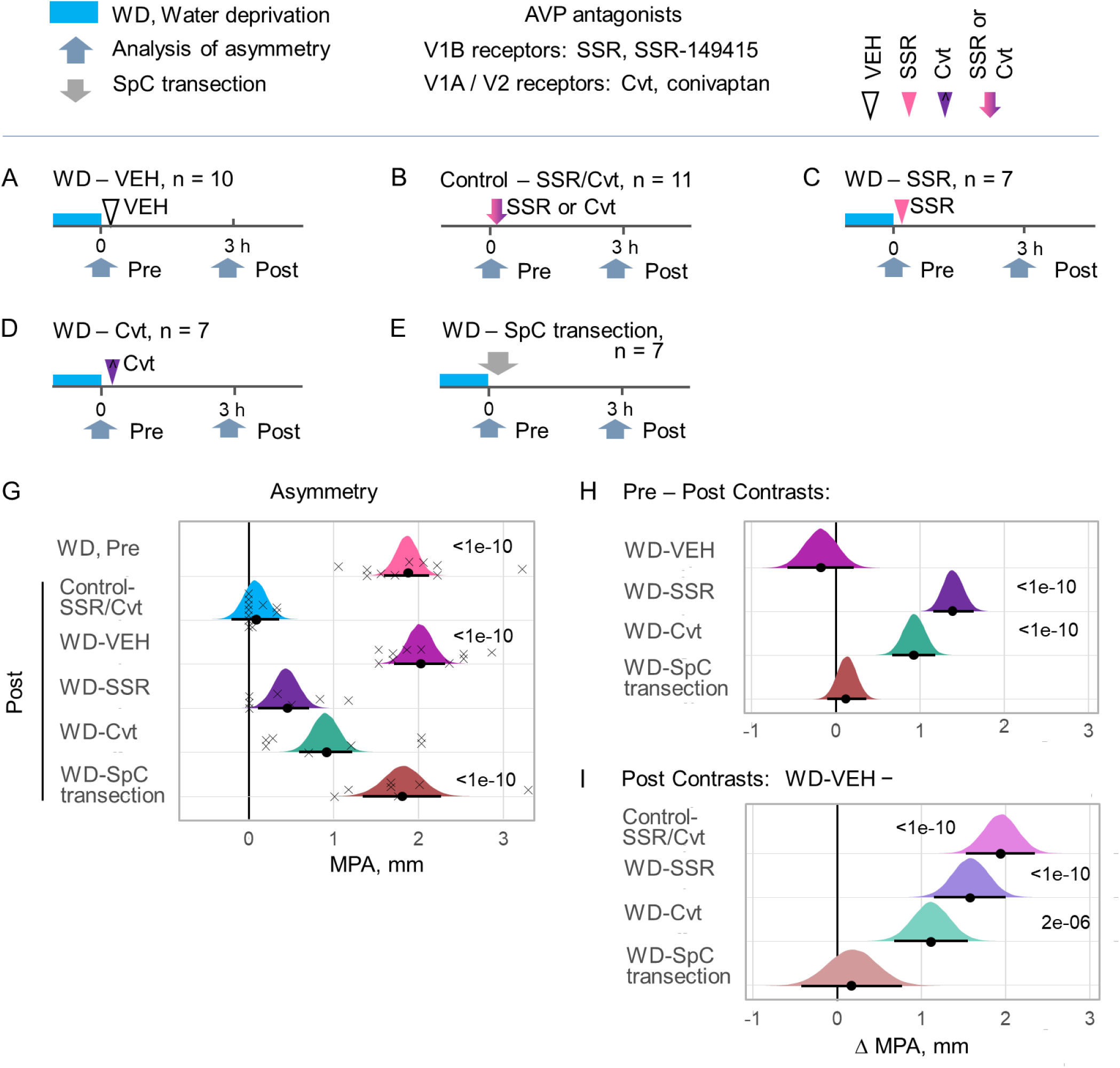
Effects of V1B and V1A/V2 receptor antagonism on water deprivation–induced HL-PA and its persistence after spinal cord transection. Water deprivation (WD) induced HL-PA, quantified as the magnitude of postural asymmetry (MPA). The effects of SSR-149415 (SSR; V1B antagonist) and conivaptan (Cvt; V1A/V2 antagonist) were tested, as well as the persistence of WD-induced HL-PA after complete spinal cord transection. **(A–E)** Experimental design for five groups comprising WD and control rats. Procedures included: (i) two assessment time points; (ii) administration of vehicle (saline with DMSO or methylcellulose–saline suspension), SSR, or Cvt; and/or (iii) complete spinal cord transection at T2–T3. In WD groups, asymmetry was first measured after 24 h WD (Pre, baseline) (A,C–E); in control rats, baseline was measured under control conditions (B). Treatments were then administered and/or the spinal cord transected, and HL-PA was re-assessed 2–3 h later (Post). **(A)** WD rats treated with vehicle: 7 rats received 2% DMSO in saline and 3 rats received 0.6% methylcellulose–saline; no differences in MPA were detected between vehicle subgroups and they were pooled. (**B)** Control rats treated with SSR-149415 (4 rats) or conivaptan (7 rats); no differences were detected and these subgroups were pooled. **(G)** MPA values for WD rats at Pre and Post time points, and for control rats at the Post time point; crosses indicate individual animals (WD rats show MPA > 1 mm). **(H)** Within-group contrasts in MPA for WD groups, defined as ΔMPA = MPA(Post) − MPA(Pre). (I) Between-group contrasts at the Post time point comparing the WD-vehicle group (reference group with robust asymmetry) to each test group, defined as ΔMPA = MPA(WD-VEH) − MPA(test group). Posterior summaries from Bayesian regression are shown as posterior medians (black circles), 95% highest posterior density (HPD) intervals (black horizontal lines), and posterior distributions (colored density plots). Effects were considered supported when the 95% HPD interval excluded zero and the corresponding multiplicity-adjusted P value was ≤ 0.05 (adjusted P values are shown). Credible intervals are the primary inferential summaries; adjusted P values are reported as supplementary model-based estimates.

**Figure 3.**
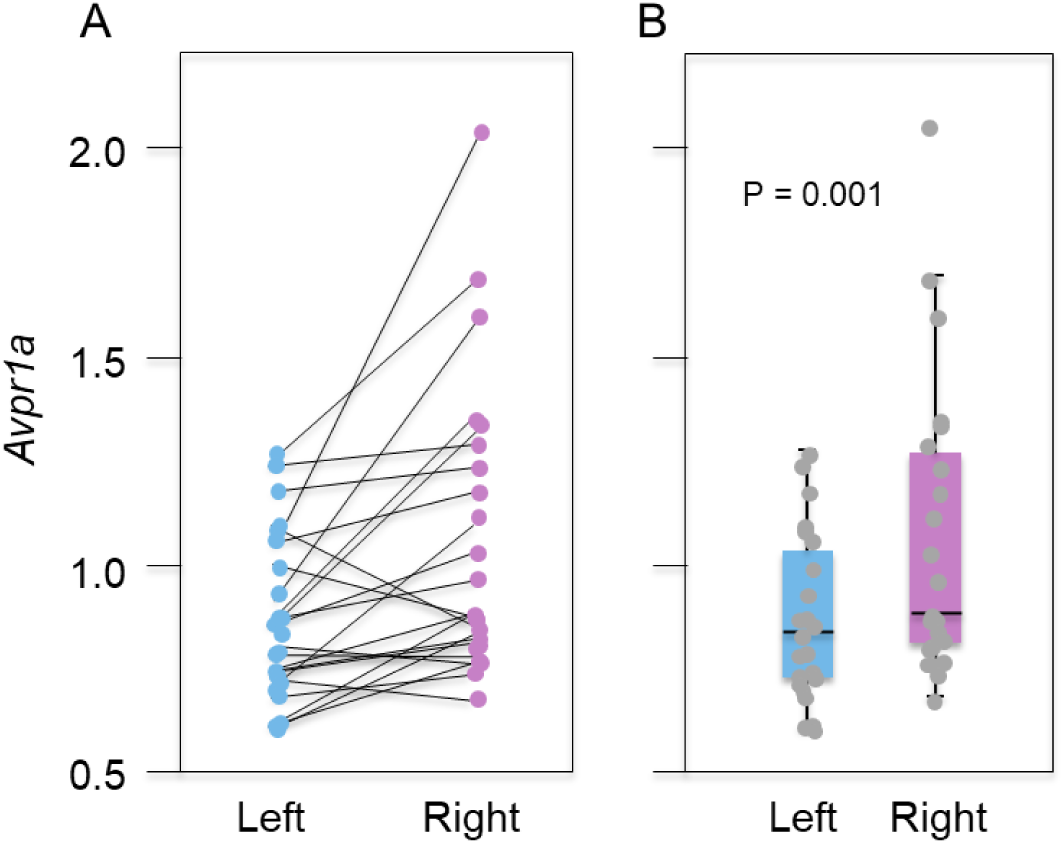
*Avpr1a* expression in the lumbar spinal cord is right-biased across animals. **(A)** Paired left–right *Avpr1a* expression values for each animal (one line per rat; n = 23). **(B)** Data are presented as boxplots with median and hinges represent the first and third quartiles. Upper and lower whiskers extend from the hinge to the highest/lowest value that lies within the 1.5 interquartile range of the hinge. mRNA levels were measured separately in left and right samples for each animal. Samples were obtained from non-related experiments with 11 sham rats and 12 rats with unilateral brain injury. No between-group difference in the asymmetry index was detected; therefore, groups were pooled for lateralization analysis. Lateralization was quantified per animal as an asymmetry index: AI_L/R_=log_2_(L/R), where (L) and (R) denote the left and right side Avpr1a mRNA levels. No between-group differences in asymmetry were revealed by comparing (AI_L/R_) and the groups were pooled. Lateralization was tested by assessing whether (AI_L/R_) differed from 0 using a two-sided Wilcoxon signed-rank test (paired, robust to non-normality): P = 0.001.

We first tested whether water deprivation (WD) can evoke hindlimb postural asymmetry (HL-PA) and whether the response is directionally biased. HL-PA was quantified using three complementary readouts (Fig. 1): magnitude of postural asymmetry (MPA), postural asymmetry score (PAS), and the proportion of asymmetric animals (Pa). PAS encodes directionality (negative values indicate left hindlimb flexion; positive values indicate right hindlimb flexion). Across rats, WD produced a strong and statistically supported HL-PA. The posterior distribution for MPA in WD rats was shifted markedly rightward relative to controls (Fig. 1A), indicating substantially larger asymmetry in WD animals (supported asymmetry, 95% HPD excluding 0; adjusted P < 1e–10). The corresponding between-group contrast (WD− control) for MPA was also positive and supported (Fig. 1B; adjusted P = 2e–07), consistent with a large WD-associated increase in asymmetry magnitude.

WD also induced a clear directional bias in the postural response. PAS values were shifted into the positive range in WD rats (Fig. 1C), indicating right hindlimb flexion as the dominant effect (supported asymmetry; adjusted P < 1e–10). The WD − control contrast in PAS was likewise supported (Fig. 1D; adjusted P = 5e–05), demonstrating that WD changes not only the magnitude but also the directionality of posture toward rightward flexion.

Finally, WD increased the proportion of animals exhibiting asymmetry (P_a_). The posterior distribution of P_a_ was close to zero in controls but markedly elevated in WD rats (Fig. 1E; supported asymmetry; adjusted P < 1e–10), and the WD − control contrast in P_a_ was strongly supported (Fig. 1F; adjusted P < 1e–10). Together, these three measures converge on the conclusion that WD induces a robust, population-level HL-PA phenotype characterized by increased asymmetry magnitude and a right-biased directional response.

### Effects of AVP receptor antagonists and spinal cord transection on WD-induced HL-PA

To probe the mechanisms underlying WD-evoked HL-PA, we next tested whether the response depends on vasopressin receptor signaling and whether it persists after disruption of descending spinal pathways. Five experimental designs were implemented (Fig. 2A–E): WD animals received vehicle (WD–VEH; n = 10), the V1B antagonist SSR-149415 (WD–SSR; n = 7), or the V1A/V2 antagonist conivaptan (WD–Cvt; n = 7); a separate WD group underwent complete spinal cord transection at T2–T3 (WD–SpC; n = 7); and control animals were tested under antagonist exposure (Control–SSR/Cvt; n = 11). HL-PA was assessed at baseline after 24 h WD (Pre) and again ∼3 h after intervention (Post).

WD animals already displayed pronounced asymmetry at baseline (Pre) (Fig. 2G; WD Pre), consistent with the robust WD phenotype in Fig. 1. In contrast, control rats (Control–SSR/Cvt) showed low MPA at the Post time point (Fig. 2G), indicating that antagonist exposure per se did not induce HL-PA under control conditions.

In vehicle-treated WD animals, MPA remained high at the Post time point (Fig. 2G; WD–VEH), showing persistence of the WD phenotype over the measurement interval. When SSR-149415, a selective V1B receptor antagonist, was administered after WD, Post MPA was markedly reduced (Fig. 2G; WD–SSR). This reduction was supported by the within-group Pre–Post contrast (Fig. 2H; supported change with adjusted P < 1e–10) and by the between-group contrast relative to WD–VEH at Post (Fig. 2I; WD–VEH − WD–SSR; supported difference with adjusted P < 1e–10). Thus, V1B signaling is required for full expression of the WD-evoked HL-PA phenotype.

Similarly, administration of conivaptan (V1A/V2 antagonist) strongly reduced WD-evoked MPA (Fig. 2G; WD–Cvt). The within-group Pre–Post contrast indicated a supported reduction (Fig. 2H; adjusted P < 1e–10), and the Post contrast relative to WD–VEH was also supported (Fig. 2I; adjusted P = 2e–06). Notably, the suppression produced by conivaptan appeared consistent with a model in which V1A/V2 signaling contributes to the expression or maintenance of asymmetry rather than being the sole required step.

Finally, WD-evoked HL-PA persisted after complete thoracic spinal cord transection. Post MPA in the transected WD group (WD–SpC transection) remained elevated and did not show a meaningful reduction relative to WD–VEH at the Post time point (Fig. 2G,I; WD–VEH − WD–SpC transection contrast centered near 0). This persistence argues that the WD-evoked asymmetry does not require intact descending spinal pathways at the time of expression and is therefore compatible with a humoral (blood-borne) component.

Together, these results show that WD produces robust HL-PA that is suppressed by V1B antagonism and reduced by V1A/V2 antagonism, while remaining expressed after complete spinal cord transection, supporting the involvement of endocrine/humoral signaling in generating the asymmetric posture.

### *Avpr1a* expression in the left and right lumbar spinal cord

Because the WD phenotype is directionally biased (right hindlimb flexion) and is sensitive to antagonism of vasopressin receptors, we examined whether components of vasopressin signaling in the spinal cord are themselves laterally organized. We measured *Avpr1a* mRNA in paired left and right lumbar spinal samples (Fig. 5; n = 23 rats).

Paired within-animal comparisons revealed a consistent rightward bias: Avpr1a expression was higher on the right than on the left in the majority of animals (Fig. 5A). This lateralization was confirmed by testing the asymmetry index (AI_L/R_=log_2_(L/R), where L and R denote the left and right side *Avpr1a* mRNA levels) against zero using a two-sided Wilcoxon signed-rank test, which supported right-biased expression (Fig. 5B; P = 0.001). Thus, *Avpr1a* expression in lumbar spinal cord is laterally organized, providing a plausible “decoder” substrate through which circulating AVP-dependent signals could be translated into side-specific motor output.

## Discussion

The first finding of this study is that WD induces HL-PA characterized by right hindlimb flexion. This effect persisted after complete spinal transection suggesting that the asymmetry is maintained through humoral signaling but not commands descending from the brain to the spinal cord through the neural tracts. WD is a symmetric homeostatic challenge that engages the lamina terminalis–hypothalamic osmoregulatory network and the AVP system, including the subfornical organ/organum vasculosum lamina terminalis-driven thirst and feedforward/feedback regulation of AVP secretion [29,30]. WD and water restriction also recruits endocrine stress-effector pathways mediated by the proopiomelanocortin-derived adrenocorticotropic hormone and β-endorphin, an endogenous opioid peptide [31,32]. In parallel, WD recruits PVN “pre-autonomic” control of sympathetic outflow via PVN projections to the spinal cord and brainstem (e.g., the rostral ventrolateral medulla), providing an anatomical route by which a symmetric internal state can be translated into coordinated neuroendocrine–autonomic output [33,34]. No evidence, as far as we know, has been presented that WD by itself can drive left- or right-side dominant PVN program. Thus, a conservative interpretation of our findings – the WD lateralization effects – is that HL-PA and its direction (right hindlimb flexion) are set downstream of a symmetric neuroendocrine activation rather than by lateralized PVN recruitment.

Our findings corroborate previous study showing that symmetric challenge – cold and immobilization stress as well as visceral pain – also produces asymmetric response–HL-PA in rats [35]. The postural effects were evident after complete spinal cord transection and left limb flexion predominated as a trend. The effects were apparently mediated via opioid receptors – the opioid antagonist naloxone blocked the HL-PA.

### The AVP HL-PA mechanism

The WD-evoked humoral mechanism may be mediated, at least in part, by neuroendocrine AVP signaling. This interpretation is supported by two observations: (i) AVP receptor antagonists abolish WD-induced HL-PA, and (ii) AVP itself, delivered symmetrically (intravenously or intracisternally), as well as chemogenetic activation of hypothalamic AVP neurons, elicits HL-PA with right hindlimb flexion [21,36]. In these experiments, the effects of exogenous AVP and AVP-neuron were observed in rats after complete spinal cord transection, indicating that they can be transmitted via humoral pathways rather than through descending neural tracts.

The finding that antagonists of both V1B and V1A/V2 receptors suppress WD-induced HL-PA is consistent with a two-site (“two-hit”) AVP mechanism, in which circulating AVP released from magnocellular hypothalamic neurons engages both a distal spinal target and a proximal pituitary target (Figure 4). In the spinal cord, AVP can enhance neuronal excitability and modulate synaptic transmission [37–39]. AVP directly depolarizes identified motoneurons and can facilitate inhibitory transmission in ventral horn neurons, effects mediated predominantly via V1A rather than V1B receptors. Consistently, AVP binding sites are present at high density in somatic motor nuclei of the spinal cord [40]. In addition, AVP signaling influences spinal sensory processing and plasticity in pain pathways [41,42], further supporting the capacity of spinal V1A signaling to reshape sensorimotor network excitability. Together, these findings provide a physiologically grounded route by which circulating AVP could bias postural output via spinal circuitry.

**Figure 4.**
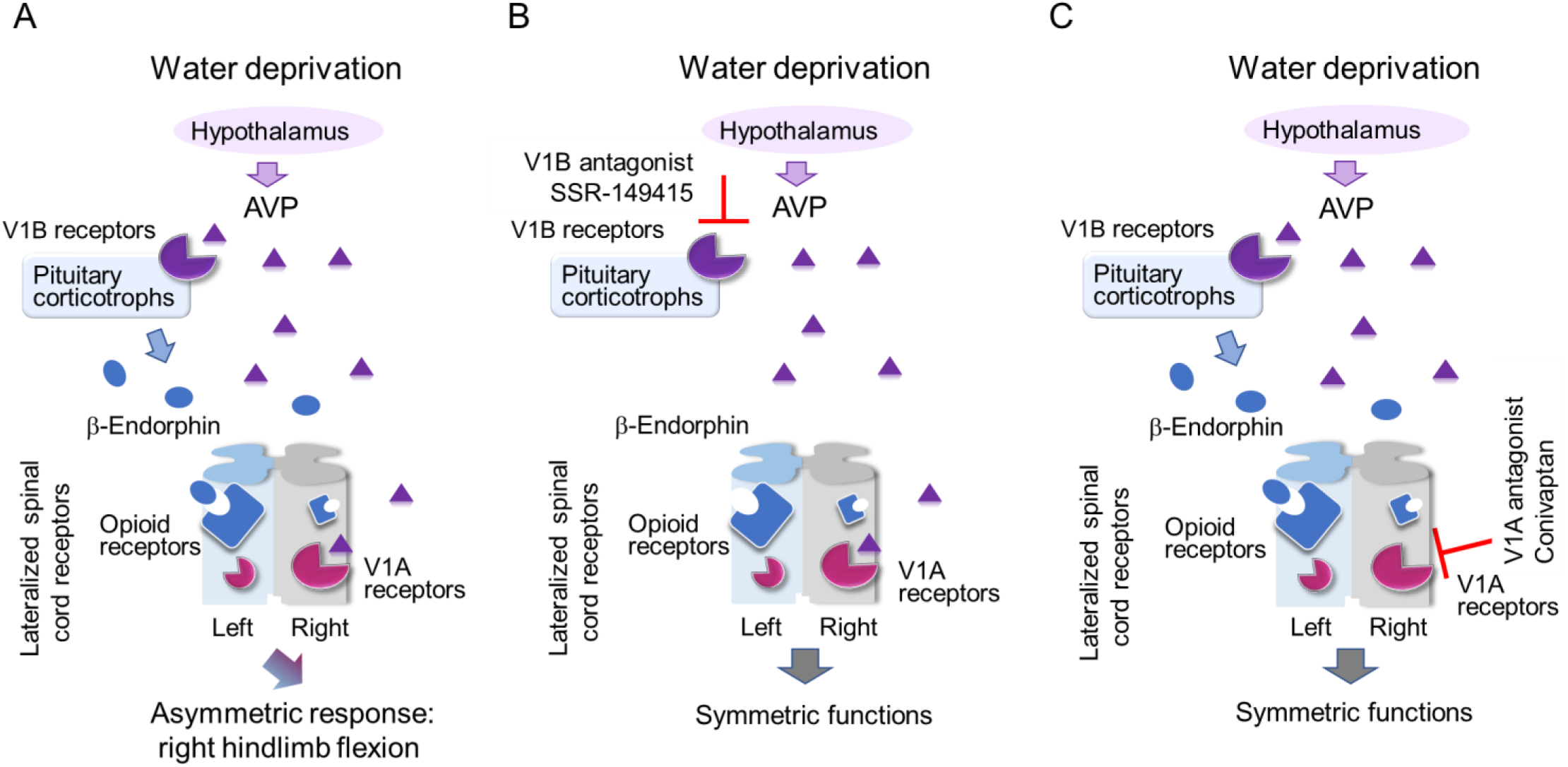
Hypothetical two-site (“two-hit”) model for water deprivation (WD)–induced HL-PA. **(A)** The observation that both the V1B antagonist SSR-149415 and the V1A/V2 antagonist conivaptan suppress HL-PA is consistent with a model in which circulating AVP released from magnocellular hypothalamic neurons engages two functionally distinct targets: (1) a proximal pituitary site that contributes to the asymmetric state via endocrine output, and (2) a distal spinal site that modulates sensorimotor excitability. In this framework, the pituitary “hit” is mediated by V1B receptors on corticotrophs, through which AVP facilitates secretion of POMC-derived hormones, including β-endorphin [43]. Circulating β-endorphin can induce HL-PA with right hindlimb flexion [21]. The spinal “hit” is mediated by V1A receptors within ventral horn sensorimotor circuitry, where AVP modulates neuronal excitability and synaptic transmission [37–39], consistent with dense vasopressin binding in spinal motor nuclei [40]. We further propose that left–right specificity is determined by lateralized “decoder” mechanisms in the spinal cord: Avpv1r (V1A receptor) expression is biased to the right spinal cord, whereas opioid receptor gene expression is biased to the left (this study and REFS). Together, these lateralized receptor systems could enable circulating AVP and β-endorphin to evoke side-specific motor output by differentially biasing mirror-symmetric spinal circuits. **(B,C)** Antagonism at either site is predicted to suppress the asymmetric state: V1B blockade would reduce AVP-driven pituitary output (including β-endorphin release), whereas V1A/V2 blockade would attenuate AVP actions on spinal sensorimotor circuitry. Consequently, asymmetric motoneuron drive is reduced and symmetry in posture and reflexes is restored. In this two-hit framework, the individual actions of AVP and POMC-derived peptides may be necessary, while their combination may be sufficient to break left–right symmetry under WD conditions.

A plausible proximal site is the pituitary gland, where V1B receptors on corticotrophs mediate AVP-potentiated release of pro-opiomelanocortin (POMC)-derived hormones, including ACTH and β-endorphin [43] (Figure 4). In turn, circulating β-endorphin can contribute to formation of right-flexion HL-PA as it was previously revealed [21]. Mechanistically, β-endorphin may act via opioid receptors expressed on spinal interneurons in dorsal and ventral horn circuits that regulate sensory processing, reflex integration, and motor output [43–46]. Opioid receptors are expressed by inhibitory interneuron classes (including premotor Ia inhibitory interneurons and Renshaw cells), and opioids modulate ventral root reflexes through both presynaptic inhibition of afferent input and postsynaptic effects on interneurons controlling motoneuron activity [44,48]. Notably, spinal motor actions of opioid peptides and opioids are preferentially directed toward pathways driven by flexor reflex afferents.

Previous studies demonstrated that HL-PA can be evoked in spinalized animals by AVP, β-endorphin, enkephalins, and synthetic opioids and their effects are side-specific: AVP and β-endorphin induce right hindlimb flexion, whereas Met-enkephalin and κ-opioid agonists (U50,488H, bremazocine, dynorphin) induce left hindlimb flexion [21–24]. This side specificity suggests that AVP and opioid peptides differentially bias mirror-symmetric spinal circuits controlling left and right hindlimb muscles, potentially through asymmetrically distributed receptors. Consistent with this idea, expression of *Avpr1a* (encoding the V1A receptor) is right-biased in spinal cord (this study). Opioid receptors are also expressed asymmetrically, with μ, δ, and κ receptor transcripts showing left-side dominance [23,49], while their relative proportions differ between spinal halves and are coordinated across dorsal and ventral regions in a side-dependent manner. Together, lateralized spinal AVP and opioid systems provide a physiologically grounded route through which circulating AVP and POMC-derived opioid peptides could produce a right-biased postural response in WD rats.

It is worth noting that AVP and opioid peptides exert side-specific postural effects in rats whose descending neural control of spinal circuits was abolished by complete transection of the spinal cords [21,23,24]. However, HL-PA with right limb flexion develops in WD rats with an intact spinal cord, suggesting that WD diminishes signaling via descending neural tracts, which would otherwise interfere with the asymmetric influence of neurohormones in naïve animals.

Our findings corroborate previous study showing that cold and immobilization stress as well as visceral pain also produce asymmetric response–HL-PA in rats [35]. The postural effects were evident after complete spinal cord transection and left limb flexion predominated as a trend. The effects were apparently mediated via opioid receptors – the opioid antagonist naloxone blocked the HL-PA.

### Limitations

This study has several limitations. First, we assessed water deprivation (WD)–evoked lateralization using the HL-PA model under anesthesia. This approach provides a robust and standardized readout of side-specific output and helps minimize supraspinal contributions, thereby isolating a neuroendocrine component. However, it does not capture the full spectrum of posture and motor coordination in awake animals. In freely behaving WD rats, voluntary and reflex postural corrections may compensate for an endocrine-driven bias, whereas reduced descending control under anesthesia may unmask it as persistent asymmetry. Future studies should therefore test whether WD induces measurable lateralized motor bias in awake animals using complementary behavioral paradigms, which would strengthen translational relevance.

Second, although WD produced lateralized effects, the underlying neuroendocrine mechanism remains incompletely resolved. The anatomical sites at which AVP initiates and maintains the response—and the principal loci of antagonist action (hypothalamic, pituitary, spinal, vascular, or peripheral)—cannot be determined from the present design.

Third, it remains unclear whether the relevant humoral signal is elevated AVP alone or a shift in the balance of circulating factors that bias left–right output, with AVP being one component (e.g., involvement of POMC-derived peptides such as β-endorphin). Our findings with exogenous AVP and chemogenetic activation of AVP neurons support a role for AVP in initiating HL-PA, while antagonist experiments indicate that AVP signaling is also required to maintain the asymmetric state, suggesting that initiation and maintenance may share overlapping neuroendocrine mechanisms. Identifying upstream events that recruit the AVP system—and disentangling initiation versus maintenance mechanisms—will require experimental strategies beyond receptor antagonism and is outside the scope of the present study.

Additional limitations include reliance on a single V1B antagonist and a single V1A/V2 antagonist. While SSR-149415 is widely used as a V1B antagonist in rodents [50–56], pharmacological specificity cannot be fully established without antagonist replication and site-restricted manipulations. Moreover, blockade by conivaptan is consistent with a spinal AVP component, but because conivaptan also targets V2 receptors and can influence systemic fluid balance, resolving V1A- versus V2-dependent contributions will require a V1A-selective antagonist and/or spinally localized manipulations.

Finally, to test the “multiple circulating signals” hypothesis directly, future work should profile the plasma neuropeptidome after WD—including AVP and POMC-derived peptides (e.g., ACTH, β-endorphin/β-LPH, and MSH)—and determine whether candidate factors exert side-specific activity. Despite these constraints, the present findings provide a foundation for mechanistic dissection of lateralized hypothalamic control of peripheral physiology at circuit and behavioral levels under physiological conditions.

### Conclusions

WD-induced HL-PA can be viewed as a functional readout of neuroendocrine symmetry breaking, in which AVP-linked hormonal states can determine a directional peripheral motor response under symmetric stimulation. WD is a spatially symmetric systemic challenge, yet it produces a population-level directional HL-PA characterized by right hindlimb flexion. Thus, WD appears to recruit an internal symmetry-breaking mechanism that converts a non-directional homeostatic state into directional postural “set-point” (“symmetric challenge → asymmetric response”). AVP signaling—acting via V1B and V1A-dependent mechanisms may initiate and maintain (‘lock’) this set-point once formed. The findings rise questions on (i) how general the phenomenon of side-specific processing of external symmetric stimuli and left-right sided response is to them, and on (ii) the side-specific neurohormonal regulation of paired organs and tissues as a driving force of such responses in the bilaterians.

## Data availability

Data supporting the findings of this study and all codes used for analysis are available within the article, its Supporting Information and on https://github.com/YaromirKo/biostatistics-nms

## Supplementary material

Supplementary material will be available online.

## Methods

### Animals

Wild type male Wistar rats were obtained from Janvier-labs, France. The animals received food and water ad libitum and were maintained in a 12-h day-night cycle (light on from 10:00 p.m. to 10:00 a.m.) at a constant environmental temperature of 21°C (humidity: 65%). Ethical approval was obtained from the Malmö/Lund ethical committee of animal experiments (Dnr. 5.8.18-17317/2021).

### Experimental design and drug treatment

The surgical procedures, and time points for treatments and physiological measurements are shown in Figures 1 and **2**. Rats were water deprived for 24 h. Subgroup of water deprived rats underwent complete spinal cord transection after 24-h water deprivation. HL-PA and stretching force were analyzed after 24 h water deprivation period (the 0-hour time point was designated as “Pre”) and then approximately 3 h later after pharmacological treatment or spinal cord transection (designated as “Post”).

SSR-149415 and conivaptan were injected immediately after the first asymmetry analysis. SSR-149415 was dissolved in 0.6% methylcellulose-saline suspension and injected intraperitoneally at 5 mg/kg. The SSR149415 dose was chosen based on previous studies, in which 1–10 mg/kg i.p. doses were observed to efficiently block endocrine processes and behaviors in a variety of models [52–56]. The dose 5 mg / kg, i.p. was used but not optimized due to its sufficiency to block the behavioral effects of AVP neuron activation. Conivaptan was diluted in 5% dimethyl sulfoxide (DMSO) and injected intravenously (i.v.) at 0.3 mg/kg. The dose was chosen based on previous studies, in which 0.3–3 mg/kg i.v. doses were observed to efficiently block endocrine processes and behaviors in a variety of models [57–59].

The design of control and water deprived rat groups exposed to the antagonists and vehicle is shown in Figure 2A-E.

### Spinal cord transection

The animals were anesthetized with isoflurane (1.5% isoflurane in a mixture of 65% nitrous oxide and 35% oxygen). Core temperature of the animals was controlled using a feedback-regulated heating system. The anaesthetized animals were mounted onto the stereotaxic frame and the skin of the back was incised along the midline at the level of the upper thoracic vertebrae. After the back muscles were retracted to the sides, a laminectomy was performed at the T2 and T3 vertebrae. A 3-4-mm spinal cord segment between the two vertebrae was dissected and removed [21,22]. The completeness of the transection was confirmed by (i) inspecting the cord during the operation to ensure that no spared spinal tissue bridged the transection site and that the rostral and caudal stumps of the spinal cord were completely retracted; and (ii) examining the spinal cord in all animals after termination of the experiment.

### Analysis of HL-PA

The HL-PA value and the side of the flexed limb were assessed as described elsewhere using the hands-off method [21,22]. Briefly, the measurements were performed under isoflurane anesthesia (1.5% isoflurane in a mixture of 65% nitrous oxide and 35% oxygen). The level of anesthesia was characterized by a barely perceptible corneal reflex and a lack of overall muscle tone. The anesthetized rat was placed in the prone position on the 1-mm grid paper.

To measure HL-PA, silk threads were glued to the nails of the middle three toes of each hindlimb, and their other ends were tied to one of two hooks attached to the movable platform that was operated by a micromanipulator constructed in the laboratory [60]. To reduce potential friction between the hindlimbs and the surface with changes in their position during stretching and after releasing them, the bench under the rat was covered with plastic sheet and the movable platform was raised up to form a 10° angle between the threads and the bench surface. The limbs were adjusted to lie symmetrically, and stretching was performed over 1.5 cm at a rate of 2 cm/sec. The threads then were relaxed, the limbs were released, and the resulting HL-PA was photographed. The procedure was repeated three times in succession, and the HL-PA values for a given rat were used in statistical analyses.

The limb that projected over a shorter distance from the trunk was considered to be flexed. The HL-PA was measured in mm with negative and positive HL-PA values that were assigned to rats with the left and right hindlimb flexion, respectively. The HL-PA was assessed by the magnitude of postural asymmetry (MPA) that shows absolute asymmetry size and by the postural asymmetry score (PAS) that shows the HL-PA value and flexion side. The MPA does not show a flexion side. These two measures are obviously dependent; however, they are not redundant and for this reason, all are required for characterization of the HL-PA data and presentation.

### Analysis of hindlimb resistance to stretch

Stretching force was analyzed under isoflurane anesthesia using the micromanipulator-controlled force meter device constructed in the laboratory [60]. Two Mark-10 digital force gauges (model M5-05, Mark-10 Corporation, USA) with a force resolution of 50 mg were fixed on a movable platform operated by a micromanipulator. Three 3-0 silk threads were glued to the nails of the middle three toes of each hindlimb, and their other ends were hooked to one of two force gauges. The flexed leg of the rat in the prone position was manually stretched to the level of the extended leg; this position was taken as 0 mm point. Then both hind limbs were stretched caudally, moving the platform by micromanipulator at a constant rate of 5 mm/sec for 10 mm. No, or very little, trunk movement was observed with stretching for the first 10 mm; therefore, the data recorded for this distance were included in statistical analysis. The forces (in grams) detected by each of the two gauges were simultaneously recorded (100 Hz frequency) during stretching. Three successive ramp-hold-return stretches were performed as technical replicates. Because the entire hindlimb was stretched, the measured resistance was characteristic of the passive musculo-articular resistance integrated for hindlimb joints and muscles [61–63]. The resistance analyzed could have both neurogenic and mechanical components, but their respective contributions were not distinguished in the experimental design. The resistance was measured as the amount of mechanical work W_L_ and W_R_ to stretch the left- and right hindlimbs, where W was stretching force integrated over the stretching distance interval from 0 to 10 mm.

### Analysis of gene expression

*Avpr1a* mRNA levels were measured separately in left and right samples for each animal (n = 23) (**Supplementary Table 1**). The tissue samples were snap frozen and stored at −80 °C until assay.

*Quantitative real-time PCR.* Total RNA was purified by using RNeasy Lipid Tissue Mini Kit (Qiagen, Valencia, CA, USA). RNA concentrations were measured with Nanodrop (Nanodrop Technologies, Wilmington, DE, USA). RNA (500 ng) was reverse-transcribed to cDNA with the cDNA iScript kit (Bio-Rad Laboratories, CA, USA) according to manufacturer’s protocol.

cDNA samples were aliquoted and stored at −20°C. cDNAs were mixed with PrimePCR™ Probe assay and iTaq Universal Probes supermix (Bio-Rad) for qPCR with a CFX384 Touch™ Real-Time PCR Detection System (Bio-Rad Laboratories, CA, USA) according to manufacturer’s instructions. TagMan assay was performed in 384-well format with TagMan probes that are listed in **Supplementary Table 2.**

All procedures were conducted strictly in accordance with the established guidelines for the qRCR based analysis of gene expression, consistent with the minimum information for publication of quantitative real-time PCR experiments guidelines [64,65]. The raw qPCR data were obtained by the CFX Maestro™ Software for CFX384 Touch™ Real-Time PCR Detection System (Bio-Rad Laboratories, CA, USA). The *Avpr1a* mRNA levels were normalized to the geometric mean of expression levels of two reference genes *Actb* and *Gapdh*. GeNorm software was used to analyze the gene expression stabilities (M value) of the ten candidate reference genes (*Actb, B2m, Gapdh, Gusb, Hprt, Pgk, Ppia, Rplpo13a, Tbp*, and *Tfrc*). The calculation of the M value was based on the pairwise variation between two reference genes. If the M value was less than 1.5, it could be considered as a suitable reference gene. The smaller the M value, the higher the stability of gene expression levels (https://genorm.cmgg.be/; [66]. The expression stability of candidate reference genes was computed for all sets of samples and identified *Actb* and *Gapdh* as the most stably expressed genes. For two regions analyzed, the gene expression stability (M values) did not exceed 0.5. The optimal number of reference genes was determined by calculating pairwise variation (V value) by geNorm program. The V value for *Actb* and *Gapdh*, the top reference genes, was 0.12 that did not exceed the 0.15 threshold demonstrating that analysis of these two genes is sufficient for normalization.

### 5.8. Statistical analysis

PCR and asymmetry data were processed and statistically analyzed after completion of the experiments by the statisticians who were not involved in their execution. No intermediate assessment was performed to avoid any bias in data acquisition. The asymmetry data were obtained by unbiased hand-off method and unbiased registration of stretching force.

### Analysis of gene expression

Lateralization was quantified per animal as an asymmetry index: AI_L/R_=log_2_(L/R), where L and R denote the left and right side *Avpr1a* mRNA levels. Samples were obtained from non-related experiments with 11 sham rats and 12 rats with unilateral brain injury. Within each group (and in the pooled sample), lateralization was tested by assessing whether the AI_L/R_ differed from 0 using a two-sided Wilcoxon signed-rank test (paired, robust to non-normality). No significant differences were revealed between the groups for each side, respectively, or for the AI_L/R_ (Welch P = 0.266, Mann–Whitney P = 0.281); therefore, groups were pooled for lateralization analysis. To directly compare expression levels between sides while accounting for within-animal pairing, we additionally fit a generalized estimating equation model on log-transformed expression values with animal identifier as the clustering variable (exchangeable working correlation). The main effect of side was significant (Right > Left; two-sided P = 0.001), whereas the group × side interaction was not significant (two-sided P = 0.231).

### Processing of physiological data

Bayesian framework. Predictors and outcomes scaled and centered before we fitted Bayesian regression models via full Bayesian framework by calling *Stan 2.21.7* [67,68] using the *brms* 2.18 interface [69]. To reduce the influence of outliers, models used Student’s *t* response distribution with identity link function unless explicitly stated otherwise. Models had no intercepts with indexing approach to predictors [70]. Default priors were provided by the *brms* according to Stan recommendations [71]. Intercepts, residual SD and group-level SD were determined\from the weakly informative prior student_t(3, 0, 10). The additional parameter ν of Student’s distribution representing the degrees of freedom was obtained from the wide gamma prior gamma(2, 0.1). Group-level effects were determined from the very weak informative prior normal(0, 10). Four MCMC chains of 40000 iterations were simulated for each model, with a warm-up of 20000 runs to ensure that effective sample size for each estimated parameter exceeded 10000 [72] producing stable estimates of 95% highest posterior density (HPD) credible intervals. MCMC diagnostics were performed according to the Stan manual. P-values, adjusted using the multivariate t distribution with the same covariance structure as the estimates, were produced by frequentist summary in *emmeans* 1.8.4-1 [73] together with the medians of the posterior distribution and 95% HPD. The asymmetry and contrast between groups were defined as significant if the corresponding 95% HPD did not include zero and the adjusted P-value was ≤ 0.05.

HL-PA. Postural asymmetry. The magnitude of postural asymmetry (MPA), postural asymmetry score (PAS), and proportion of animals with postural asymmetry (P_a_) were inferred via Bayesian framework using Gaussian response distribution. Adjusted P-values for MPA ≠ 0 and PAS ≠ 0, and for the contrasts (ΔMPA, ΔPAS and ΔPa) were shown.

Stretching force. The amount of mechanical work W_L_ and W_R_ to stretch the left- and right hind limbs, respectively, was computed by integrating the smoothed stretching force measurements over stretching distance from 0 to 10 mm using loess smoothing computed by *loess* function from R package *stats* with parameters span=0.4 and family=”symmetric”. Asymmetry was assessed both as the left / right asymmetry index AI_L/R_ = log_2_ (W_L_ / W_R_). The AI_L/R_ was inferred via Bayesian framework by fitting linear multilevel models as the factor of interest. No differences in AI_L/R_ were observed between male and female subgroups, allowing pooled analysis.

## Supplementary material

**Supplementary Table 1.** The number of rats used in the study.

| Figure 1 groups | n |
| --- | --- |
| Control | 12 |
| Water-deprived | 43 |

| Figure 2 groups | Description | n |
| --- | --- | --- |
| WD-VEH | Vehicle after WD (7 DMSO-saline + 3 methylcellulose-saline; pooled) | 10 |
| Control-<br>(SSR/Cvt) | Control rats treated with SSR-149415 (4) or conivaptan (7); pooled | 11 |
| WD-SSR | WD rats treated with SSR-149415 | 7 |
| WD-Cvt | WD rats treated with conivaptan | 7 |
| WD-SpC | WD rats with complete T2-T3 transection | 7 |

| Figure 3 | n |
| --- | --- |
| Pooled (all animals) | 23 |
| Sham | 11 |
| Unilateral brain injury | 12 |

**Supplementary Table 2.** PCR probes (Bio-Rad Laboratories, CA, USA) used for analysis of the *Avpr1a* gene expression.

| Gene symbol | Gene name | Assay ID | Channel |
| --- | --- | --- | --- |
| *Actb* | *Actin beta* | qRnoCIP0050804 | FAM |
| *Avpr1a* | *Arginine Vasopressin* | qRnoCEP0023750 | FAM |
| *Gapdh* | *Glyceraldehyde-3-phosphate dehydrogenase* | qRnoCIP0050838 | FAM |

## Key resources table

| Reagent type (species) or resource | Designation | Source or reference | Identifiers | Additional information |
| --- | --- | --- | --- | --- |
| Strain, Strain background (*Rattus norvegicus*) | Wistar Hannover rat, males | Janvier-labs, France |  |  |
| Chemical compound, drug | Isoflurane | Abbott Laboratories, Norway | NDC 0044-5260-03 | Anesthesia agent |
| Commercial Assay or kit | RNasy Plus Mini kit | Qiagen, CA, USA | Cat#: 74136 | Total RNA extraction |
| Commercial Assay or kit | iScript™ cDNA Synthesis Kit | Bio-Rad Laboratories, USA | Cat#: 1708891 | cDNA Synthesis |
| Commercial Assay or kit | iTaq Universal Probes supermix | Bio-Rad Laboratories, USA | Cat# 1725131 | Real-Time PCR reagent |
| Software, algorithm | Stan | (Carpenter et al., 2017) | v 2.19.2 | Scaling the data |
| Software, algorithm | R program | (R Core Team, 2018) | v 3.6.1 |  |
| Software, algorithm | *Brms* | (Burkner, 2017) | v 2.9.6 | Interface for R |
| Software, algorithm | *Emmeans* | (Searle et al., 2012). | v 1.4 | R package |
| Other | TissuDura | Baxter, Germany | Cat#: 0600096 | Covering material |
| Chemical compound | SSR-149415 | Tocris, UK | Cat#: 6195 | Antagonist of AVP V1B receptors |
| Chemical compound | dimethyl sulfoxide (DMSO) | Tocris, UK? | Cat#: 3176? | Solvent for SSR-149415 |
| Chemical compound | Conivaptan hydrochloride | Tocris, UK | Cat#: 6830 | Antagonist of AVP V1A / V2 receptors |
| Chemical compound | Methylcellulose | SigmaAldrich | Cat#: M7027 | Solvent for SSR-149415 |

